# Testing the Vogt-Bailey Index using task-based fMRI across pulse sequence protocols

**DOI:** 10.1101/2025.02.06.636866

**Authors:** Kristian Galea, Aitor Alberdi Escudero, Nina Attard-Montalto, Nolan Vella, Robert E. Smith, Maria Mora Corral, Christine Farrugia, Paola Galdi, Kenneth Scerri, Liam Butler, Claude J. Bajada

## Abstract

Local connectivity analyses in fMRI such as the Vogt-Bailey Index, investigate the prevalence of co-fluctuations in the time-series of adjacent voxels. While there have been *in silico* assessments of the VB Index, this technique has not yet been assessed systematically. This study has two aims: first, to assess the VB Index using a task paradigm with well established *a priori* expectations on the brain region predominantly responsible for executing this task to determine whether the VB Index specifically highlights this area. Second, we investigate if, and how, the spatial resolution of the sequence protocols employed, with their inherent effects on the signal-to-noise ratio, affect the resultant VB maps. A cohort of 10 research volunteers under-went fMRI acquisitions utilising a block design finger tapping experiment. Each volunteer was scanned with three sequence protocols, with all parameters equivalent except for the volume of the voxels. Task-associated VB maps derived from statistical comparison of normal VB maps with task-regressed VB maps were compared with those obtained from the conventional General Linear Model approach. Particular emphasis was placed on the identification of the hand portion of the motor homunculus. Across sequence protocols, the VB Index consistently identified elevated local connectivity in, qualitatively, the same portion of the motor cortex as that yielded by the General Linear Model based on the task paradigm. The consistently significant task-associated increases in the VB Index, across sequence protocols, in the cortical region associated with an fMRI task paradigm, provides support for the biological relevance of such local connectivity measures.

## Introduction

A subset of the various data analysis methods employed in functional Magnetic Resonance Imaging (fMRI) are local connectivity methods. These data analysis methods are predicated on the assumption that the voxels encompassed by a brain region with elevated neural activity are temporally similar to each other, reflecting the underlying synchronous neural activity.^1, 2^ Therefore, local connectivity methods infer brain function by assessing the similarity between the time-series signal of local voxel neighbourhoods. One such local connectivity measure of relevance to this work is the VB Index,^3^ a technique based on spectral graph theory that employs a searchlight approach to iteratively assess local connectivity of voxel neighbourhoods.

A desirable attribute of local connectivity methods is that they do not require explicit prior knowledge on what the participant is doing, cognitively while they are lying in the scanner. This makes local connectivity methods particularly suited for resting-state fMRI data analysis where prior knowledge may not be readily available.^1^ However, an inherent challenge with resting-state fMRI analysis is the lack of readily available “ground truth”. Thus, assessing local connectivity methods *in vivo*, particularly in resting-state fMRI is challenging.

One possible way to perform *in vivo* assessment of local connectivity methods would be to evaluate their outcomes in an experiment where activity of a known brain region is explicitly evoked and verifiable using alternative analysis techniques. This is the definition of task-based fMRI, where there is a rich history of task paradigms and association of brain regions to such.^4–7^ Typically, in task-based fMRI, each image voxel is analysed using the General Linear Model (GLM),^8^ which models the voxels time-series signal as a weighted sum of regressors, including the Blood Oxygenation Level Dependent (BOLD) signal predicted by explicit engagement and disengagement with the task. Statistical tests are then applied to identify voxels with a time-series that exhibits a significant fit to the model.^9–11^ Any meaningful local connectivity algorithm should ideally exhibit elevated values in the brain regions with association with the task performed. Although such assessment has been previously performed for an earlier local connectivity measure, Regional Homogeneity (ReHo),^12, 13^ the VB Index has thus far only been validated using synthetic data.^14^

Therefore, this study aims to provide a systematic assessment of the VB Index using such a well-established task. We investigate whether the VB Index identifies the primary region of interest for an associated task. Specifically, in this study, the task performed is that of finger tapping, and the associated primary region of interest is the motor cortex. To provide a reference map for this assessment, we employ a standard GLM analysis on the same data.^15^

Given the mechanism by which the VB Index is calculated, it was hypothesized that the spatial resolution of the fMRI acquisition may have a considerable impact on its ability to identify task-related activation where the functional region is physically small. For conventional GLM analysis, it is only necessary to measure at least one image voxel with a time series correlated with the task. With adequate signal-to-noise ratio (SNR), the model will suitably associate that voxel with the task. Further, fMRI data are typically smoothed prior to model fitting, sacrificing spatial specificity for sensitivity to the effect. For the VB Index to become elevated however requires that multiple adjacent image voxels co-fluctuate with one another, hence only being sensitive to functional regions that encompass multiple image voxels. Relatedly, larger voxels and hence poorer resolution may encompass an entire functionally-homogeneous region within a single voxel, rendering the VB index ineffective at detecting local connectivity. Yet if the spatial resolution is too high the sensitivity of the VB Index to that co-fluctuation may be corrupted by low SNR.^16^ Further, performing smoothing of fMRI data is expected *a priori* to artificially elevate the VB Index everywhere given that it would explicitly impose co-fluctuations in adjacent voxels. Given these confounds, our investigation of the VB Index was performed at multiple acquisition resolutions, and with two alternative strategies for data smoothing.

## Methods

### Participants

Ten research volunteers (aged 22 to 55 years; mean±standard deviation 31.5±10.4 years) were recruited for the study. Nine of the volunteers were male and one was female. Seven of the volunteers were right-handed. Prior to the MRI scanning, written informed consent was obtained from the participants. In order to be eligible to participate, the volunteers had to have normal or corrected-to-normal vision. The study was granted ethics approval by the Faculty of Health Science Research Ethics Committee (FREC) at the University of Malta. This study was conducted in accordance with relevant guidelines and regulations, including the EU’s General Data Protection Regulation (GDPR) for data privacy, and the Declaration of Helsinki for ethical principles regarding medical research involving human subjects.

### Scanning protocols

A SIEMENS 3T MAGNETOM Vida (Siemens Healthcare GmbH, Erlangen, Germany) MRI system was used throughout. The BioMatrix Head/Neck 64 channel coil (Siemens Healthcare GmbH, Erlangen, Germany) was used for reception of the radiofrequency signal during the MRI scanning. The volunteers were positioned in a supine position on the MRI gantry, with their head at the centre of the head coil. Each volunteer was scanned for a period of approximately one hour.

Each volunteer was scanned using a high-resolution T1-weighted MPRAGE axial pulse sequence (1^3^ mm^3^ isotropic voxels, TE=2.66ms, TR=2190ms, TI=925ms, flip angle=8^◦^, GRAPPA; R=2). Then, a gradient field map image was acquired (2.5^3^ mm^3^ isotropic voxels, TR=529ms, TE_1_=4.92ms, TE_2_=7.38ms, flip angle=60^◦^). Eight of the ten volunteers were scanned with a high-resolution T2-weighted SPACE axial sequence (1^3^ mm^3^ isotropic voxels, TE=417ms, TR=3140ms, variable flip angle, GRAPPA; R=2, turbo factor = 282).

The volunteers were scanned using three different gradient echo planar imaging (EPI) pulse sequences. The three EPI sequences, referred to as the 2.5^3^ mm^3^, 2.0^3^ mm^3^ and 1.8^3^ mm^3^ sequences, employed consistent scanning parameters (flip angle=65^◦^, TR=1840ms, TE=34ms, simultaneous multi slice (SMS) with a multiband factor (MB) of 4, phase partial Fourier of 6/8) except for the volume of the voxels which was set to 2.5^3^ mm^3^ (field of view (FOV)=240×240mm^2^, in-plane resolution=96×96, Bandwidth=2084Hz/Px), 2^3^mm^3^ (FOV=240×240mm^2^, in-plane resolution=120×120, Bandwidth=2084Hz/Px), or 1.8^3^ mm^3^ (FOV=241×241 mm^2^, in-plane resolution=134×134, Bandwidth=2072Hz/Px). For each EPI sequence, a total of 60 axial slices (with no slice gaps) and 393 volumes were acquired, with an acquisition time of approximately 12 minutes. The order in which the different sequences were acquired for each participant was shuffled.

### fMRI experimental paradigm

Participants were required to complete a finger tapping task while being scanned with the three EPI sequences. The task required participants to tap either their right thumb or index finger at the onset of a particular visual stimulus. This was done using the NordicNeuroLab response grip (NordicNeuroLab AS, Bergen, Norway). The experimental task was explained to participants before scanning commenced.

The experimental finger tapping task was designed using PsychoPy.^17^ The task instructions were synchronised with the fMRI volume acquisition using the NordicNeuroLab SyncBox (Nordic-NeuroLab AS, Bergen, Norway). The times at which the buttons were pressed and the times at which the fMRI volumes were acquired were stored.

The task included three visual stimuli, a white fixation cross shown during rest periods at the centre of the screen, and the active stimuli were presented as coloured crosses on the right side of the screen, with a red cross corresponding to tapping of the right thumb and blue cross corresponding to tapping of the right index finger. All three stimuli were presented on a solid grey background. Participants were instructed to tap their right thumb at the onset of the red cross, and to tap their right index finger at the onset of the blue cross. Participants were required to repeatedly tap either the thumb or the index finger until the end of each trial, at which point the white fixation cross was presented, upon which they were instructed to rest and stop tapping their thumb/finger. Each experimental trial lasted about 18.4s (10 fMRI volumes), and was followed by a rest period of equal length. For each block within the set of experimental trials, the tapping finger (thumb or index) was chosen at random, but the overall structure of alternating task (tapping) and rest periods remained consistent throughout the scan. Each participant was presented with 20 task blocks.

### Data pre-processing

The MRI data were first structured according to the brain imaging data structure (BIDS) standard^18^ using the “BIDSKIT” tool (https://github.com/jmtyszka/bidskit). fMRIPrep (version 23.2.0)^19, 20^ was then used to preprocess the structural and functional data. The recommended fMRIPrep boilerplate text is reproduced verbatim in the next few subsections except for minor adjustments for formatting, language, and clarity.

### Preprocessing of B_0_ inhomogeneity mappings

A total of 10 fieldmaps were found available within the input BIDS structure. Then, for each volunteer, a B0 nonuniformity map (or fieldmap) was estimated from the phase-drift maps measured with two consecutive gradient-recalled echo (GRE) acquisitions. The corresponding phase-maps were phase-unwrapped with ‘prelude’ (FSL version 6.0^21–23^).

### Anatomical data preprocessing

A total of 10 T1-weighted (T1w) images were available within the input BIDS dataset. Each T1w image was corrected for intensity non-uniformity (INU) with ‘N4BiasFieldCorrection‘,^24^ distributed with ANTs 2.5.0,^25^ and used as T1w-reference throughout the workflow. The T1w-reference was then skull-stripped with a Nipype implementation of the ‘antsBrainExtraction.sh’ workflow (from ANTs), using OASIS30ANTs as target template. Brain tissue segmentation of cerebrospinal fluid (CSF), white-matter (WM) and gray-matter (GM) was performed on the brain-extracted T1w using ‘fast’ [FSL (version 6.0),^21–23]^. Brain surfaces were reconstructed using ‘recon-all’ [FreeSurfer 7.3.2,^26^], and the brain mask estimated previously was refined with a custom variation of the method to reconcile ANTs-derived and FreeSurfer-derived segmentations of the cortical gray-matter of Mindboggle.^27^

Where a T2-weighted image was available, it was used to improve pial surface refinement. Brain surfaces were reconstructed using ‘recon-all’ [FreeSurfer 7.3.2,,^26^ and the brain mask estimated previously was refined with a custom variation of the method to reconcile ANTs-derived and FreeSurfer-derived segmentations of the cortical gray-matter of Mindboggle.^27^ Volume-based spatial normalisation to one standard space (MNI152NLin2009cAsym) was performed through nonlinear registration with ‘antsRegistration’ (ANTs 2.5.0,^25, 28^), using brain-extracted versions of both T1w reference and the T1w template. The following template was selected for spatial normalization and accessed with TemplateFlow [23.1.0,:^29^ ICBM 152 Nonlinear Asymmetrical template version 2009c [;^30, 31^ TemplateFlow ID: MNI152NLin2009cAsym].

### Functional data preprocessing

For each of the 3 BOLD runs found per subject, the following preprocessing was performed. First, a reference volume was generated, using a custom methodology of fMRIPrep, for use in head motion correction. Head-motion parameters with respect to the BOLD reference (as rigid-body transformation matrices) were estimated using ‘mcflirt’ [FSL 6.0,^32^] before any spatiotemporal filtering. The estimated fieldmap was then aligned with rigid-registration to the target EPI (echo-planar imaging) reference run. The field coefficients were mapped to the reference EPI using the transform. The BOLD reference was then co-registered to the T1w reference using ‘bbregister’ (FreeSurfer) which implements boundary-based registration.^33^ Co-registration was configured with six degrees of freedom. Several confounding time-series were calculated based on the preprocessed BOLD: framewise displacement (FD), framewise displacement and the derivative of the root mean squared variance over voxels (DVARS), and three region-wise global signals. FD was computed using two formulations following Power et al., (2012) (absolute sum of relative motions,^34^) and Jenkinson et al., (2012)^32^ (relative root mean square displacement between affines). FD and DVARS were calculated for each functional run, both using their implementations in Nipype [following the definitions by Power et al., (2012)^34^]. Three global signals were extracted within the CSF, the WM, and the whole-brain masks. Additionally, a set of physiological regressors were extracted to allow for component-based noise correction [CompCor,^35^]. Principal components were estimated after high-pass filtering the preprocessed BOLD time-series (using a discrete cosine filter with 128s cut-off) for the two CompCor variants: temporal (tCompCor) and anatomical (aCompCor). tCompCor components were then calculated from the top 2% variable voxels within the brain mask. For aCompCor, three probabilistic masks (CSF, WM and combined CSF+WM) were generated in anatomical space. The implementation differs from that of Behzadi et al., (2007)^35^ in that instead of eroding the masks by 2 pixels in BOLD space, a mask of pixels that likely contain a volume fraction of GM is subtracted from the aCompCor masks. This mask is obtained by dilating a GM mask extracted from FreeSurfer’s aseg segmentation, and it ensures components are not extracted from voxels containing a minimal fraction of GM. Finally, these masks were resampled into BOLD space and binarized by thresholding at 0.99 (as in the original implementation). Components were calculated separately within the WM and CSF masks. For each CompCor decomposition, the k components with the largest singular values were retained, such that the retained components’ time-series were sufficient to explain 50 percent of variance across the nuisance mask (CSF, WM, combined, or temporal). The remaining components were dropped from consideration. The head-motion estimates calculated in the correction step were also placed within the corresponding confounds file. The confound time-series derived from head motion estimates and global signals were expanded with the inclusion of temporal derivatives and quadratic terms for each 38. Frames that exceeded a threshold of 0.5 mm FD or 1.5 standardized DVARS were annotated as motion outliers. Additional nuisance time-series were calculated by means of principal component analysis of the signal found within a thin band (crown) of voxels around the edge of the brain.^36^ All resamplings were performed with a single interpolation step by composing all the pertinent transformations (i.e. head-motion transform matrices, susceptibility distortion correction when available, and co-registrations to anatomical and output spaces). Gridded (volumetric) resamplings were performed using ‘nitransforms‘, configured with cubic B-spline interpolation.

Many internal operations of fMRIPrep use Nilearn 0.10.2 [http://nilearn.github.io/], mostly within the functional processing workflow. For more details of the pipeline, see the section corresponding to workflows in fMRIPrep’s documentation (https://fmriprep.readthedocs.io/en/latest/workflows.html).

### Data analysis

#### General Linear Model Analysis

The preprocessed fMRI data for each volunteer acquired with each EPI sequence was analysed using the GLM. The GLM was constructed using the Python library Nilearn (version 0.10.3). The GLM analysis consisted of a first-level GLM computed for each subject separately. The second-level GLM which provided statistical inferences at a group level was performed using the “Randomise” tool of the FMRIB Software Library (FSL).

Prior to the first-level statistical testing, the fMRI data of each subject were spatially smoothed to improve the SNR and account for inter-subject anatomical variability.^37^ A Gaussian kernel with a full width at half maximum (FWHM) of 4.806 mm was selected based on the recommendations of Weibull et al., (2008)^38^ for voxels of volume 1.8^3^ mm^3^. To ensure consistency, the same FWHM was applied for all image series regardless of acquired resolution.

Three mass-univariate first-level GLMs were fit to the smoothed data from each volunteer (one per sequence protocol). For each first-level GLM, the design matrix contained a single experimental task regressor that modelled the BOLD signal response during the thumb and index tapping task period. The experimental regressor was modelled as the convolution of the Glover Haemodynamic Response Function (HRF)^39^ and the stimulus function of the experiment. In this case, the stimulus function had a duration equivalent to one stimulus period, i.e., 10 fMRI volumes; 10×1.84=18.4s. The timings of the on-off period used to construct the boxcar stimulus function were acquired from the NordicNeuroLab system. The design matrices also contained a number of confound regressors, extracted by fMRIPrep, that modelled various nuisance parameters including 24 motion-related regressors. A total of 10 discrete cosine transform regressors generated by fMRIPrep were also included in the design matrix to model low-frequency physiological and scanner noise. The first 6 principal components, extracted by fMRIPrep using principal component analysis (PCA) on regions of interest identified by the anatomical CompCor method as being unlikely to be modulated by neural activity^20, 35^ were also included in the design matrix for each participant. Finally, the global signal regressor was also included in the design matrix since it were shown to improve confound regression performance when used with CompCor.^40^ Outlier fMRI volumes containing sudden and large motion artefacts were scrubbed from the GLM analysis, by applying thresholds to the DVARS determined by fMRIPrep for each fMRI volume in each run. Thresholds of 0.5 mm for the FD,^41^ and 2 for the standardised DVARS^34^ were chosen.

The GLM model was then fit to the spatially smoothed fMRI data for each volunteer and sequence separately. The effect size corresponding to the task regressor was then computed.

The MNI152NLin2009cAsym T1-weighted image with a resolution of 2.0^3^ mm^3^ was chosen as the primary common template, which was then downsampled to 3.0^3^ mm^3^ resolution (such that transforming processed data to template space would involve downsampling only) using trilinear interpolation with FSL.^21^ The resulting statistical maps were spatially normalised to the template image during the preprocessing stage using the ANTs library.^28^

A second-level GLM analysis was performed per acquisition resolution for group-level inferences using a mass-univariate one-sample t-test on the spatially normalised statistical maps. The “Randomise” tool of FSL^42^ was used for the second-level analysis. Statistical maps were thresholded with an alpha level of 0.05 (p<0.05) using threshold-free cluster enhancement (TFCE) family-wise error rate (FWER) correction.

#### The Vogt-Bailey Index

This study aimed to assess the ability of the VB Index to highlight regions associated with the motor task. To achieve this aim, the VB Index maps were compared against corresponding VB maps computed on data with the modelled task-component subtracted. Therefore, the task-related component of the fMRI time series was first modelled using the GLM. The fitted task model was subtracted from the time-series. Both the normal data and task-regressed data were then passed through a denoising pipeline prior to VB Index computation. This is because while the GLM permits addition of confounding signals to the model itself for regression of their explanatory power, calculation of the VB Index offers no such mechanism. Therefore, prior to the VB analysis, a denoising pipeline based on the work of Satterthwaite et al., (2013)^41^ was applied to the fMRI data using the rsDenoise library^43^ (https://github.com/adolphslab/rsDenoise.git).

The preprocessed and denoised normal and task-regressed fMRI data were analysed with the VB Index. The VB Index algorithm iteratively goes over each voxel in the image, considers the voxel in the centre of the searchlight and its’ nearest neighbours, and for each neighbourhood formulates a feature similarity graph with adjacent voxels within that neighbourhood assigned as vertices on the graph connected together via edges. The weighting (*w_ij_*) assigned to each edge reflects the pairwise similarity between the time-series signal of the vertex pair (*i*, *j*) and is defined using a rectified angular similarity derived from the Pearson correlation coefficient (Equation 1).

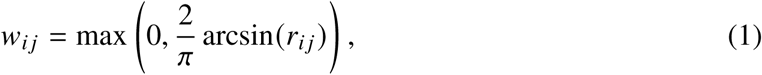

VB neighbourhoods thus consist of 7 voxels - the voxel at the centre of the searchlight and it’s nearest neighbours. The relative similarity between vertices of a graph of a single VB Index searchlight containing 7 voxels can be expressed via the similarity matrix 2.

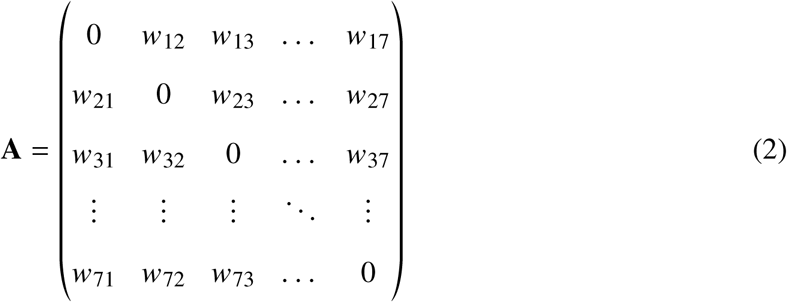

Apart from the similarity matrix, the algorithm derives the diagonal degree matrix (**D**), whose entries contain the sum of edge weights incident on each vertex. The unnormalised graph Laplacian is then defined as (**L** = **D** − **A**). Partitioning the graph according to the RatioCut criterion is approximated using a spectral relaxation, whose solution is determined by the eigenvector associated with the second-smallest eigenvalue of *L*. This eigenvalue, *λ*_2_, is known as the algebraic connectivity of the graph. In the VB Index, *λ*_2_ is normalised by the number of vertices *n* in the neighbourhood. The final normalised metric is termed as the VB Index (Equation 3).

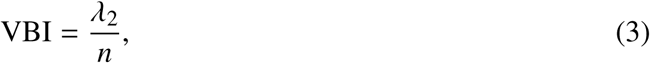

Higher VB Index values indicate stronger correlation within the time-series signal of the local neighbourhood; more specifically, the higher the VB Index, the less readily can the graph be disconnected into two parts (as such, the VB Index behaves somewhat as an edge detection technique).^3, 14^ Conversely, smaller VB Index values indicate smaller correlations in the time-series signal of the voxel neighbourhood. The algorithm iteratively performs this procedure for each voxel in the image.

Both normal and task-regressed time-series data were analysed with the VB Index using two approaches. The VB method 1 (VB1) replicated the smoothing operation performed during the GLM analysis for fair comparison. Therefore, in VB1 the denoised data were first smoothed with a Gaussian FWHM of 4.806 mm prior to the computation of the volumetric VB Index. In the VB method 2 (VB2), the volumetric VB Index was computed directly from the denoised data. Figure 1 illustrates the workflow of both pipelines.

**Figure 1:**
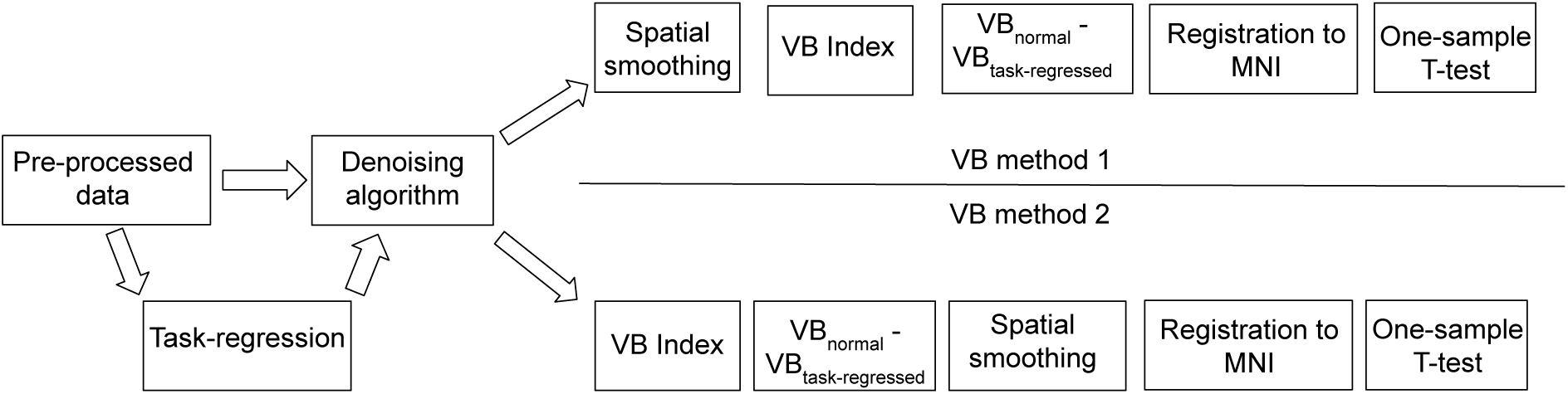
The workflow of the VB method 1 and VB method 2.

The VB Index calculated from the original time-series was then compared with that calculated from the task-regressed time series to quantify the change in local connectivity between the normal VB maps and the task-regressed VB maps. The VB Index computed on the task-regressed time-series data was subtracted from the original VB Index maps. For the VB2 approach, the difference maps were spatially smoothed to account for residual inter-subject anatomical variability. Subject-level difference maps were then spatially normalised to the 2.0^3^ mm^3^ MNI152NLin2009cAsym template downsampled to 3.0^3^ mm^3^, as was done during the GLM analysis. The data was grouped and a one-sample t-test applied using the non-parametric “Randomise” tool of FSL^42^ to assess whether the group mean differs from zero. Statistical maps were thresholded with an alpha level of 0.05 (p<0.05) using TFCE FWE correction. These maps will be hereafter be referred to as task-association VB maps.

#### Comparison of statistical maps

The GLM and task-associated VB maps were compared across sequence protocols to assess the consistency of results across sequence protocols. This was achieved using the Dice-Sørensen coefficient (DC) which is defined as the ratio of the intersection between a pair of statistical maps (*X* and *Y*) to the to the average number of statistically significant voxels in both statistical maps as shown by equation 4.^44^ This procedure was performed for both the statistical maps obtained with the GLM and the VB Index. Cross-comparison between the task-associated VB maps and GLM were also performed using the Dice coefficient.

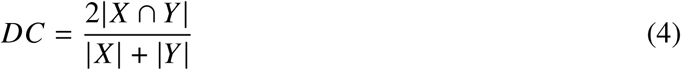

Finally, cross-comparison of the GLM and task-associated VB maps were also performed using the Overlap coefficient (OC). The Overlap coefficient assesses the ratio of the intersection between the two statistical maps to the total number of statistically significant voxels in the smaller statistical map as given in equation 5.^45^ Thus, the Overlap coefficient determined the proportion to which the smaller statistical map was enveloped within a larger statistical map.

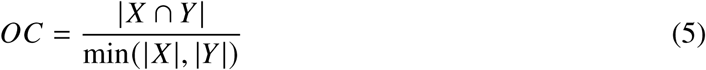

## Results

Three separate analysis pipelines were run on the preprocessed fMRI data; two different pipelines of the VB Index, and a classical GLM analysis. The VB1 and VB2 pipelines were performed after subtracting task-regressed VB maps from the VB original maps to specifically query local functional connectivity of task-related co-fluctuations. The first VB approach (hereafter VB1) replicated the smoothing operation commonly adopted for conventional GLM analysis, where the fMRI data are spatially smoothed prior to model fitting to increase the SNR. The second VB approach (hereafter VB2) involved the application of spatial smoothing after the VB analysis. In VB2 smoothing was carried out to account for residual misalignment in functional localisation relative to anatomy. Further, the order of smoothing for VB2 conformed with the standard VB Index pipeline of computing the metric on unsmoothed data. Given that the VB Index is an edge-detection technique sensitive to functional boundaries,^14^ it was hypothesized that operating on unsmoothed data would better localise the contrast of functional boundaries. Images will always be displayed in radiological view.

### GLM results

Slices from the 1-TFCE-corrected-p value GLM statistical maps obtained for each sequence protocol are given in Figure 2. Activation was detected consistently across sequence protocols contralaterally to the task performed in the precentral gyrus of the motor cortex, an Omega shaped region in the axial plane, commonly referred to as the “hand knob” region. Furthermore, activity was detected consistently contralaterally in the thalamus, clearly seen from the coronal slice in Figures 2a, 2b and 2c. Cerebellar activity was detected predominantly ipsilaterally, whilst occipital activity was detected predominantly contralaterally to the task performed.

**Figure 2:**
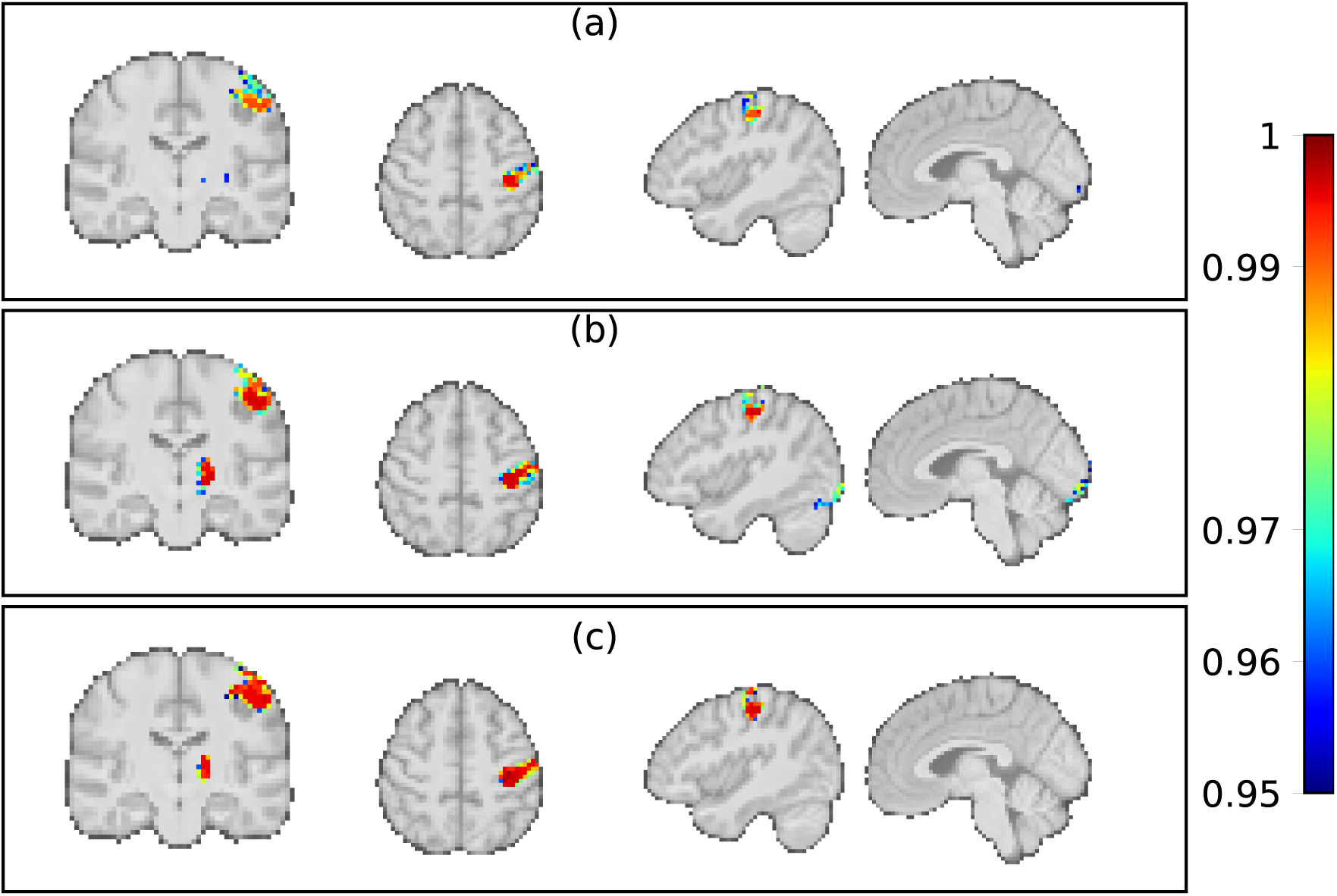
TFCE thresholded (p<0.05) GLM statistical maps. Colours indicate 1-TFCE-corrected-p values. The second-level statistical maps are overlaid on the group average T1-weighted image spatially normalised to the MNI152NLin2009cAsym template. The first row (a), second row (b) and third row (c) displays the t-maps from the 1.8^3^ mm^3^, 2.0^3^ mm^3^, and 2.5^3^ mm^3^ sequences respectively.

### VB Index results

Figures 3 and 4 show slices from the resulting 1-TFCE-corrected-p value statistical task-associated maps obtained using VB1 and VB2 respectively. A significant positive difference indicate that the modelled task-related component contributes to the elevated VB Index in that region.

**Figure 3:**
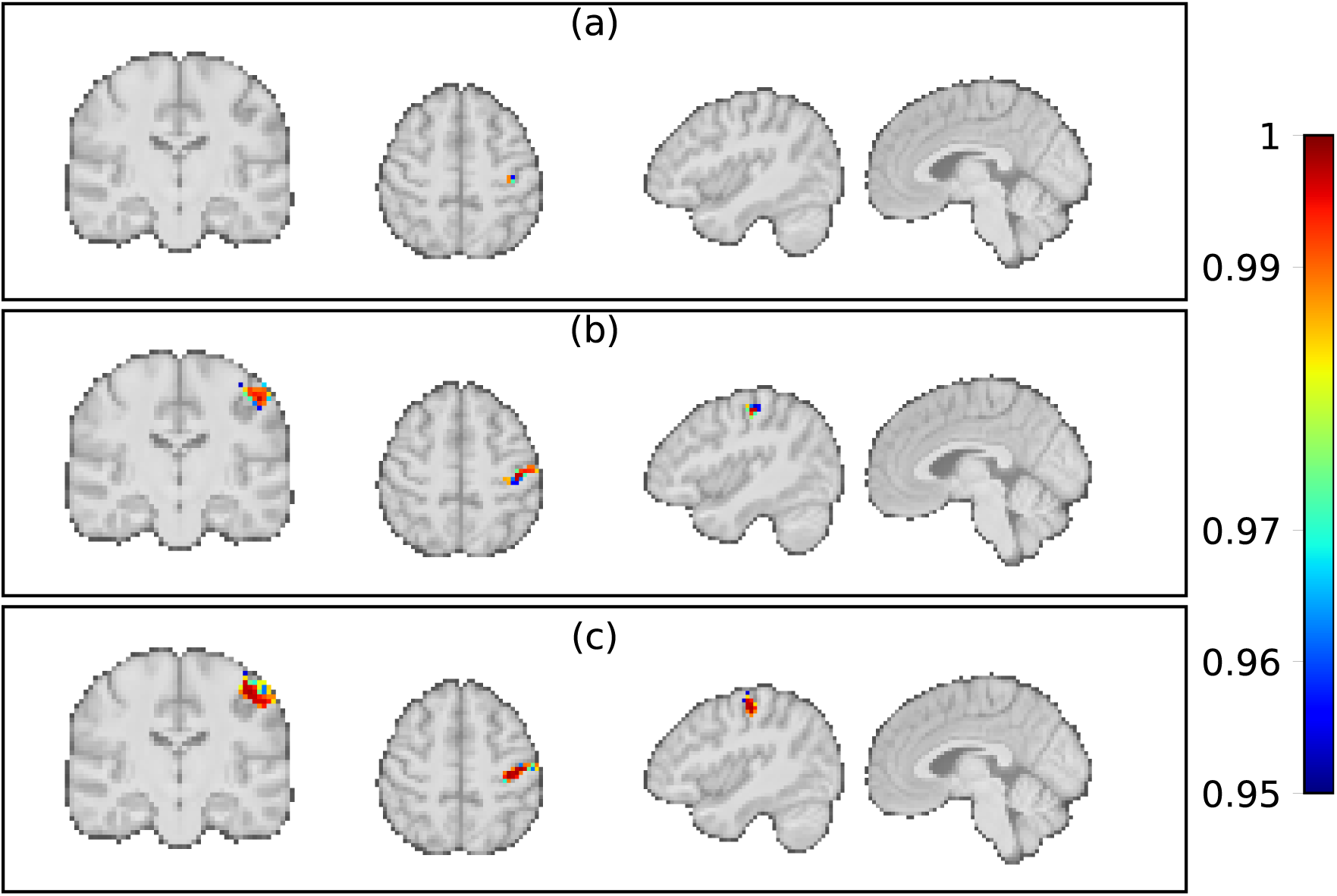
Task-associated TFCE thresholded (p<0.05) statistical maps obtained with VB1. Colours indicate 1-TFCE-corrected-p values. The first row (a), second row (b), and third row (c) display the statistical maps obtained with the 1.8^3^ mm^3^, 2.0^3^ mm^3^, and 2.5^3^ mm^3^ sequences respectively.

**Figure 4:**
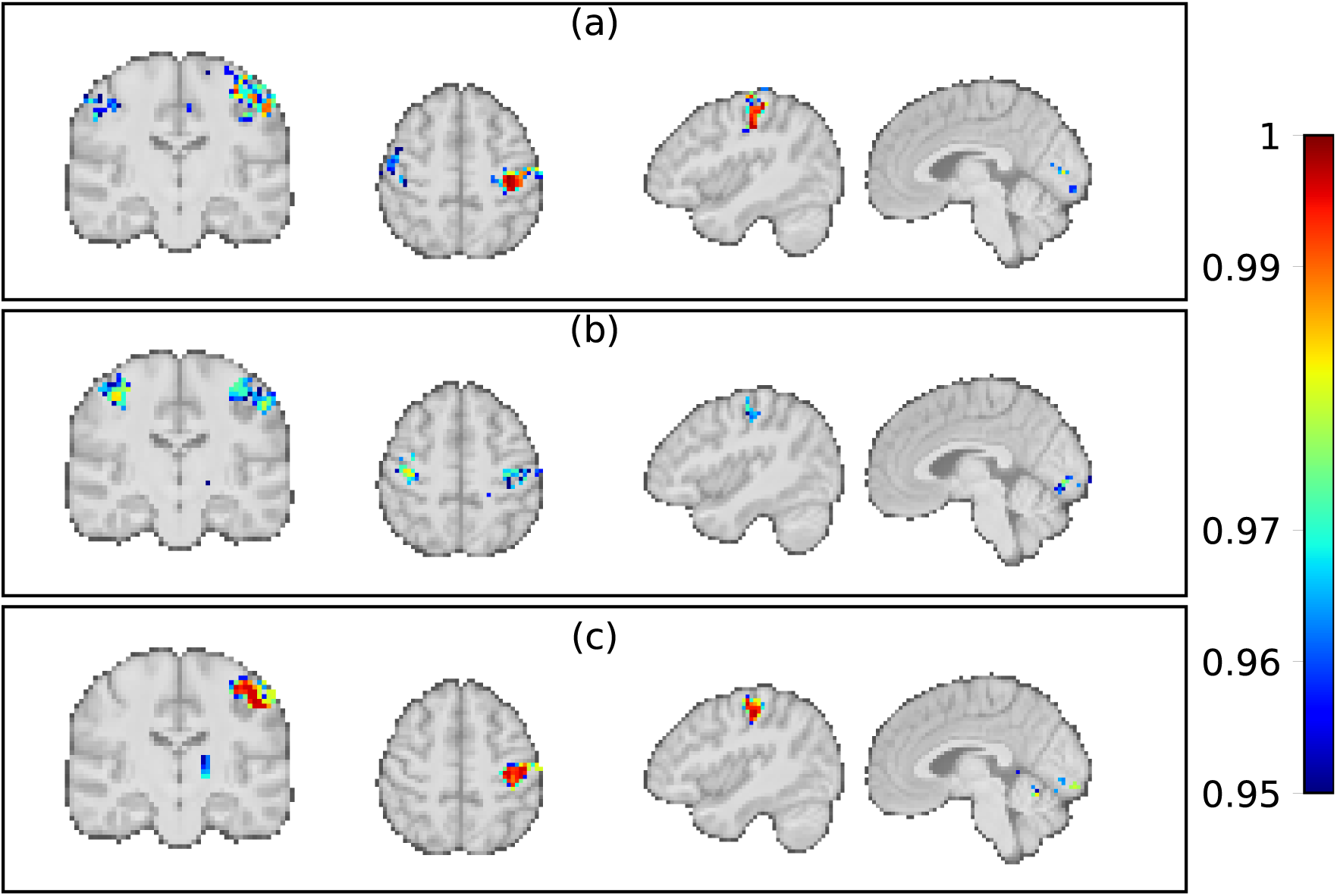
Task-associated TFCE thresholded (p<0.05) statistical maps obtained with VB2. Colours indicate 1-TFCE-corrected-p values values. The first row (a), second row (b), and third row (c) display the statistical maps obtained with the 1.8^3^ mm^3^, 2.0^3^ mm^3^, and 2.5^3^ mm^3^ sequences respectively.

In both VB pipelines, significant task-associated increases in the VB Index were found consistently across sequence protocols in the precentral gyrus of the motor cortex in agreement with the GLM (Figure 2). Task-associated VB1 maps highlighted connectivity predominantly in the precentral gyrus contralaterally to the task performed with some ipsilateral connectivity in the anterior cerebellar lobes at the 2^3^ mm^3^ and 2.5^3^ mm^3^ voxel volume protocols. Task-associated VB2 maps also highlighted connectivity contralaterally in the precentral gyrus and cerebellum. However, VB2 maps were more expansive and also highlighted connectivity contralaterally in the thalamus at the 2^3^ mm^3^ and 2.5^3^ mm^3^ voxel volume protocols, and connectivity in the occipital lobe. The VB2 maps obtained with 1.8^3^ mm^3^ and 2.0^3^ mm^3^ voxel volume protocols also included contributions from motor areas located ipsilaterally to the task performed.

### Quantitative analysis

Given that the same task paradigm was performed by the volunteers for each sequence, it was hypothesised that the results acquired with the three sequence protocols would be consistent. Therefore, the consistency of the statistical maps across resolutions was assessed using the Dice coefficient (Figure 5). Following the nomenclature of Wilson et al., (2017),^46^ the spatial similarity between pairs of statistical maps which depends on the magnitude of the Dice coefficient shall be described as being; low if the similarity metric ranges from 0.00 to 0.19, low-moderate; 0.20 to 0.39, moderate: 0.40 to 0.59, moderate-high; 0.60 to 0.79 or high; 0.80 to 1.00.

**Figure 5:**
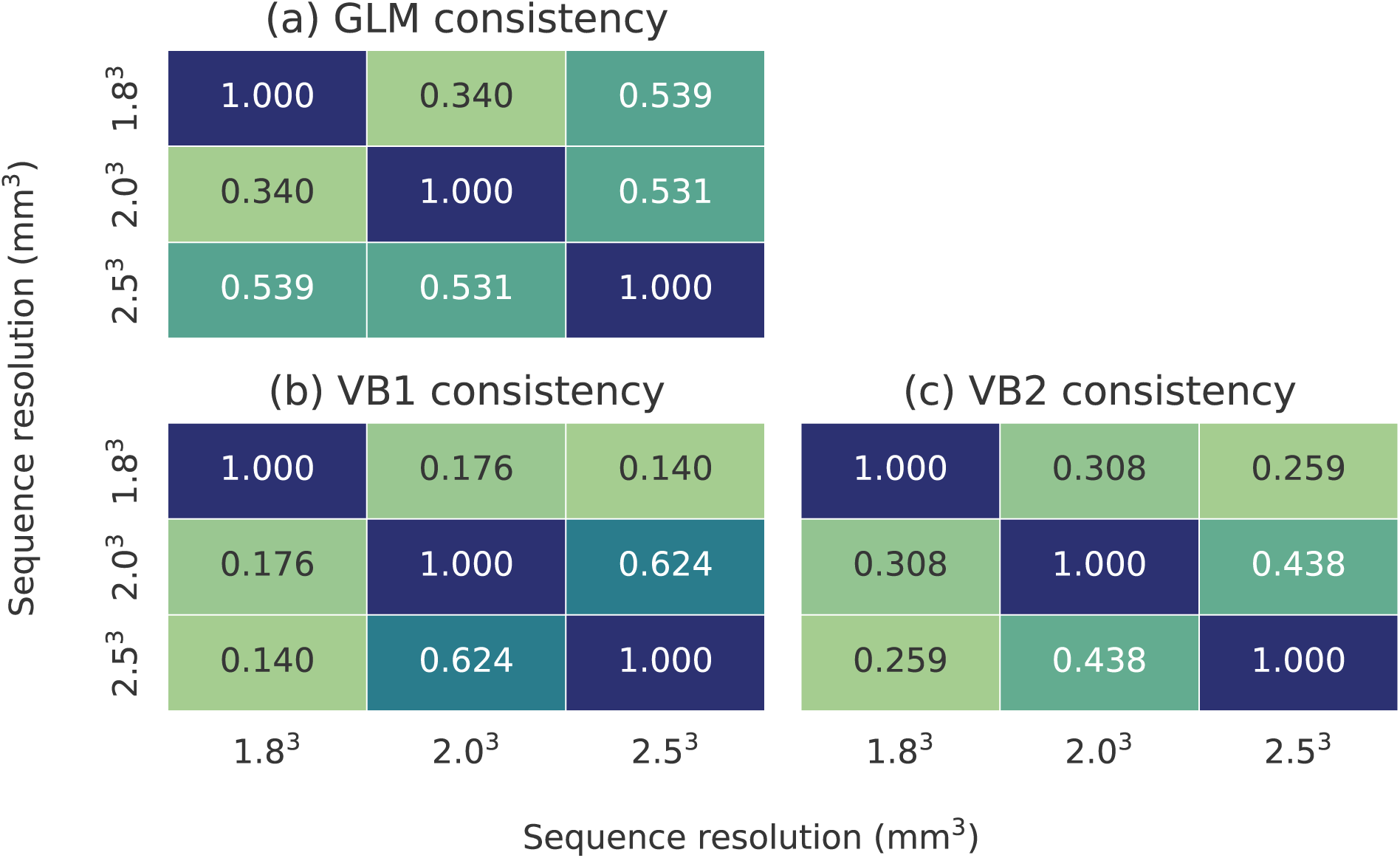
The Dice coefficients showing the relative consistency between the (a) GLM and task-associated a) VB1, b) VB2 across sequences.

The results obtained with the GLM and VB1 exhibited the highest overall consistency across pulse sequences (Figure 5a & Figure 5b), with low-moderately to moderate Dice coefficients ranging from 0.340 to 0.539 for the GLM and low to moderate-high Dice coefficients ranging from 0.140 to 0.624 for VB1. The task-associated VB2 maps were the least consistent across sequences (Figure 5c), with low-moderate to moderate Dice coefficients ranging from 0.259 to 0.438.

In order to assess the VB Index, the resulting task-associated VB maps obtained with both VB pipelines were compared to those of the GLM, a standard technique in fMRI analysis.^9^ Dice similarity between GLM and VB1 were in the low to moderate range (0.019-0.408) as seen from Figure 6. Dice scores were particularly low in VB1 for the higher 1.8^3^ mm^3^ resolution sequence. In most instances, VB2 Dice scores were higher (0.190-0.449).

**Figure 6:**
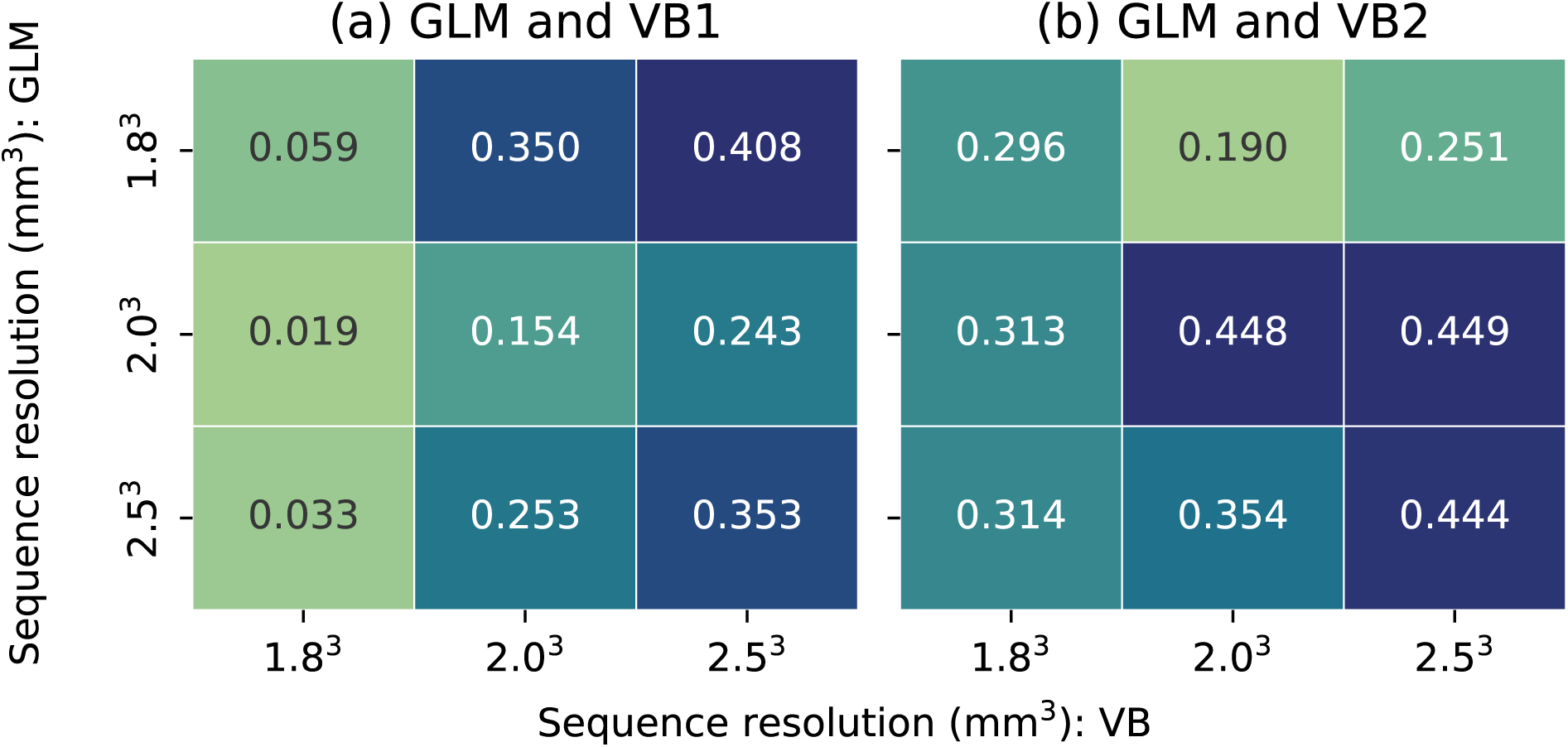
The Dice coefficients showing the relative similarity between the GLM and task-associated a) VB1, b) VB2 across sequences.

Overlap scores (Figure 7) quantify the proportion to which the smaller statistical map is contained into the larger map. The size of the statistical maps are quantified in Figure 8 by the number of statistically significant voxels per map. Figure 8 highlighted that overall VB1 maps were smaller when compared to both the VB2 and GLM which were similar in size. The size of the VB1 maps were more affected by the resolution of the sequence protocol when compared to VB1 and GLM whose size were consistent across sequence protocols. The VB1 maps had the highest overlap relative to GLM with moderate-high to high Overlap coefficients (0.600 to 0.936), implying that a substantial portion of the the smaller VB1 maps were enveloped by the GLM. VB2 maps exhibited less overlap (0.350 to 0.620) possibly due to the more expansive nature of the VB2 maps which included ipsilateral areas not present in the GLM maps. This penalised the Overlap coefficient.

**Figure 7:**
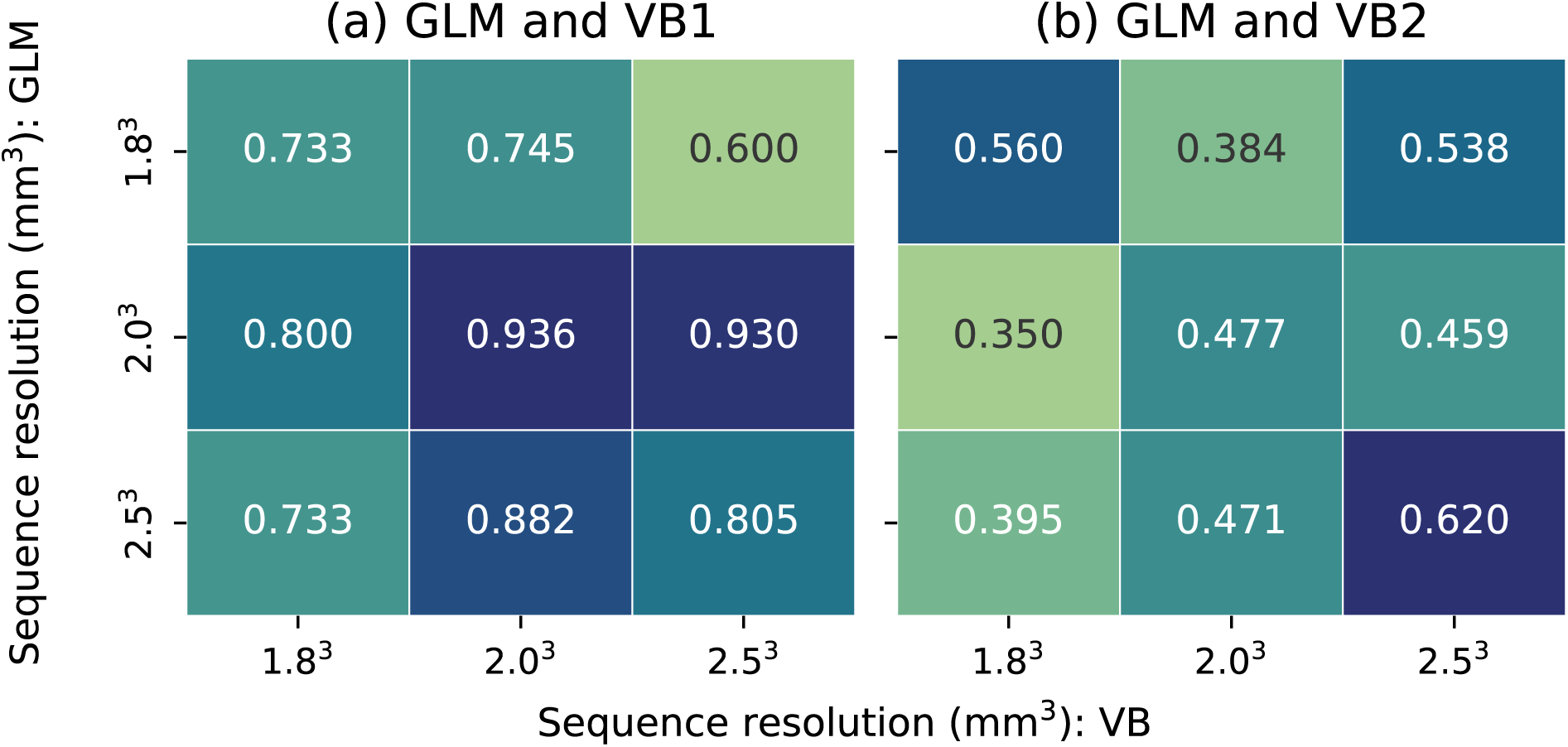
The Overlap coefficients showing the overlap between the GLM and task-associated a) VB1, b) VB2 across sequences.

**Figure 8:**
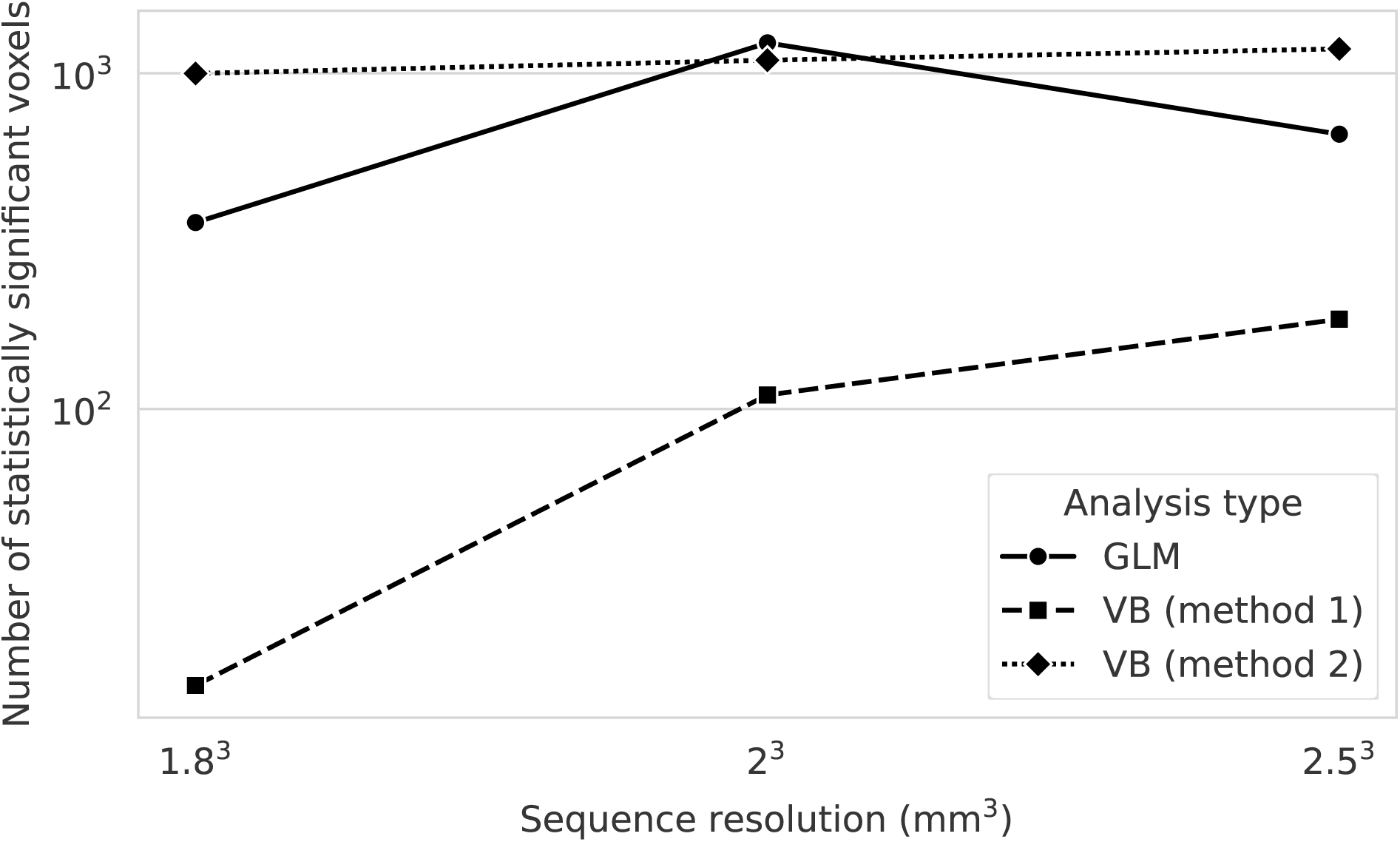
The number of statistically significant voxels highlighted after TFCE correction for the GLM and task-associated VB Index maps.

## Discussion

The lack of available ground truth in resting-state fMRI analysis makes it inherently difficult to assess local connectivity approaches. Though the VB Index was previously tested with synthetic data,^14^ it had not yet been tested on empirical fMRI data. Therefore, the study aimed to use a well established task-based experimental paradigm with a well known pattern of expected activity to assess the VB Index and its ability to specifically detect and characterise the task-related activity. Further, this assessment was carried out across sequence protocols with differing spatial resolutions and two different smoothing strategies to investigate their impact on the VB Index’s results. Results demonstrated that the VB Index consistently identified elevated local connectivity in expected task-related regions across resolutions, with clear qualitative overlap with statistical maps generated by the GLM.

The GLM analysis, revealed significant activation in the left motor cortex, contralateral to the hand used in the task, consistent with previous research.^4–7^ Activity was detected consistently in the precentral gyrus which is heavily involved in hand motor function,^47^ and contralaterally in the occipital lobe (Figures 2) which is primarily responsible for visual perception.^48^ Given that visual stimulus presentation differed subtly during task and rest conditions (central white fixation cross versus laterised coloured targets shown to the participants during the experiment), some activity in the occipital region was expected. Ipsilateral activity in the anterior cerebellar lobes and contralateral thalamic activity were also consistently highlighted in line with literature.^4–7^

The second portion of the analysis featured fitting a GLM model to the data and regressing the task block out of the data. The normal and task-regressed data were analysed using the VB Index with two separate approaches, and then compared statistically. The resulting task-associated statistical maps obtained (Figure 3 & Figure 4) in general highlighted brain areas conformal with the motor task at hand - the precentral gyrus, thalamus, occipital lobe, cerebellum. The application of smoothing before the VB analysis (VB1) produced maps predominantly confined to the precentral gyrus, whilst the application of spatial smoothing post-VB (VB2) produced maps that highlighted alternate areas such as the cerebellum, occipital lobe, thalamus (for poorer resolution), and also motor areas located ipsilateral to the task performed.

In both VB pipelines, clusters of significantly high local connectivity were identified across sequences in the precentral gyrus (Figures 3 and 4 for VB1 and VB2, respectively). The elevated local connectivity detected in the precentral gyrus with VB2 approach largely overlapped with the area identified by the GLM, while the elevated local connectivity detected in the same area with VB1 was more spatially confined, delineating a smaller region within the precentral gyrus. VB2 revealed elevated local connectivity in the occipital lobe across sequences, as seen in the right sagittal slices in Figure 4. VB2 also identified regions of elevated local connectivity in the occipital lobe, thalamus, and cerebellum conformal to the GLM, and with literature.^4, 6^ These results provide evidence that the VB Index is sensitive to local BOLD co-fluctuations associated with the experimental task.

Compared to VB2, VB1 results were predominantly confined to the precentral gyrus (see Figure 3, & Figure 4). Specifically for VB1, the number of significant voxels reduced as resolution improved (Figure 8) as evident by the smaller area of highlighted connectivity in the VB1 map (Figure 3). Conversely, resolution had little impact on the size of the VB2 map (4). Overall, smoothing prior to VB computation produced smaller task-associated VB maps which may indicate reduced sensitivity to task-evoked BOLD signal co-fluctuations. These findings suggest that the VB2 pipeline may be preferred for task-based data particularly when using higher resolution data. The findings demonstrate that the VB Index can consistently detect elevated local connectivity in brain regions with anticipated elevated task-related BOLD activity. This partially validates the VB Index’s sensitivity to coherency in the BOLD signal in empirical data.

The Dice coefficient was used for the pairwise comparison of the statistical maps generated by the different analysis methods across sequence protocols. This analysis revealed varying degrees of consistency. The GLM and VB1 results showed the highest consistency, with VB2 exhibiting the lowest consistency across sequences. In general, the statistical maps obtained with the 2^3^ mm^3^ and 2.5^3^ mm^3^ voxel volumes protocols exhibited the highest consistency. A certain degree of consistency was indeed expected across sequences as participants were subjected to the same tasks and fixed real space distance smoothing for each sequence protocol. However, perfect congruence was not anticipated due to differences in the SNR across sequences, and inherent variability in task performance which could have contributed to differences between runs.

VB2 pipeline exhibited the highest spatial similarity relative to GLM (Figure 6). The lower sensitivities of the 1.8^3^mm^3^ protocols penalised the Dice score in VB1. Although VB2 generally demonstrates greater Dice correspondence with the GLM in the task-associated analysis, VB1 appears to produce more spatially restricted changes that are predominantly centred on the expected motor hand region, particularly for the 2.0^3^ mm^3^ and 2.5^3^ mm^3^ protocols. The Overlap coefficient results (Figure 7) obtained when comparing the task-associated VB1 and GLM were, in general, moderate-high to high, signifying that in most cases, a substantial portion of the smaller (Figure 8) VB1 maps were enveloped within the larger GLM maps. The task-associated VB2 exhibited less overlap with the GLM possibly because VB2 highlighted ipsilateral motor areas which were not included in the GLM. Indeed, the task-associated VB2 and GLM maps were almost the same size (Figure 8) but the observed differences such as ipsilateral connectivity highlighted in VB2 penalised the Overlap coefficient. These results suggest that the VB Index is sensitive to task-evoked connectivity, and this connectivity exhibits strong overlap with a conventional GLM approach.

The VB results obtained at poorer resolutions had overall a higher spatial overlap to the GLM approach as they had smoother and more diffused patterns. Nevertheless, such lower resolutions may not well capture spatially small activations. Therefore, the spatial resolution should be sufficiently high to sample an activated functional area ensuring that a voxel’s neighborhood encompasses a functionally homogeneous region. However, increasing spatial resolution comes at the cost of reduced SNR which could consequently degrade the sensitivity of VB Index to the BOLD signal as was seen in VB1 (Figure 3). Therefore, optimising data acquisition for the purpose of VB analysis involves a trade-off between the competing demands of spatial resolution and SNR.

Future work to our study may focus on addressing both the limited number of participants and test paradigms used. Nevertheless, finger tapping was deliberately selected because it produces a robust and well-characterised motor cortex response. The robust nature of the expected activation patterns associated with this simple motor task was determined to be sufficient for this preliminary validation study, despite the relatively small sample size. Further work could also focus on the development of test paradigms specific for the use of local connectivity analysis for task-based fMRI data. Most of the subjects that participated in this study were male (9 male and 1 female), and future work should focus on balancing the sample to suppress gender-related imbalances. Future work could focus on the acquisition of separate resting-state data and performing a direct comparison between task-based VB maps and resting-state VB maps which would subsequently further help in clarifying the extent to which the observed maps reflect intrinsic local connectivity rather than task-related fluctuations without needing to resort to a task model as was performed in this work.

Pre-smoothing the fMRI data prior to VB Index quantification (VB1) resulted in patterns of significantly elevated local connectivity predominantly confined to the precentral gyrus, and reasonable consistency across sequence protocols with reduced sensitivity at higher resolution. VB1 maps were almost wholly enveloped by the GLM exhibiting moderate-to-high overlap with GLM results. Conversely, VB2 revealed broader patterns of significantly elevated local connectivity in expected motor areas, consistently across protocols. These findings highlight the VB Index’s sensitivity to local BOLD signal coherence and serve as preliminary validation of the VB Index. Thus, they provide evidence of its’ sensitivity to known biological effects and support further investigation in its’ application for resting state fMRI analysis. Finally, while more investigation is needed, it also suggests that the VB Index may have some utility in analysing task-based data.

## Code availability

The code used during the study is available on Github: https://github.com/KristianGalea/VB-Pulse-Sequence-Evaluation.git. Data for this study cannot be openly shared due to data protection requirements. However, the entire dataset and any derivative including scripts that were used for this article are available on the University of Malta’s data repository (drUM) in an embargoed state: https://doi.org/10.60809/drum.33077819. Any data requests can be forwarded to the University of Malta’s Magnetic Resonance Imaging Platform (UMRI) on.

## Acknowledgements

The authors would like to thank the University of Malta’s MRI Platform (UMRI) for access to scanning equipment and services, use of its infrastructure, and resources such as documentation and preprocessing scripts. Furthermore, the authors would also like to acknowledge that the study is part of the Enhancing Astronaut Neuro-Imaging Capabilities: Toolbox Optimization & Modification (Operation TOM) Project, financed by Xjenza Malta through the Space Upstream Programme 2023 (grant no. SUP-2023-01). RS is supported by fellowship funding from the National Imaging Facility (NIF), an Australian Government National Collaborative Research Infrastructure Strategy (NCRIS) capability.

## Author contributions statement

K.G: Formal analysis, Writing - original draft, Writing - review & editing. A.A.E: Formal analysis, Writing - review & editing. N.A.M: Writing - review & editing. N.V: Writing - review & editing. R.E.S: Writing - review & editing, Supervision. M.M.C: Data curation, Writing – review & editing. C.F: Writing - review & editing, Supervision. P.G: Writing - original draft, Writing - review & editing, Supervision. K.S: Writing - review & editing, Supervision. L.B: Writing - original draft, Writing - review & editing. C.J.B: Conceptualization, Writing - original draft, Writing - review & editing, Supervision, Funding Acquisition.

## References

[1] Wu Tao, Long Xiangyu, Zang Yufeng, et al. Regional homogeneity changes in patients with Parkinson’s disease Human Brain Mapping. 2009;30:1502–1510.

[2] Xu Zhe, Lai Jianbo, Zhang Haorong, et al. Regional homogeneity and functional connectivity analysis of resting-state magnetic resonance in patients with bipolar II disorder Medicine. 2019;98:e17962.

[3] Bajada Claude J., Costa Campos Lucas Q., Caspers Svenja, et al. A tutorial and tool for exploring feature similarity gradients with MRI data NeuroImage. 2020;221:117140.

[4] Gountouna Viktoria-Eleni, Job Dominic E., McIntosh Andrew M., et al. Functional Magnetic Resonance Imaging (fMRI) reproducibility and variance components across visits and scanning sites with a finger tapping task NeuroImage. 2010;49:552–560.

[5] Witt Suzanne T., Laird Angela R., Meyerand M. Elizabeth. Functional neuroimaging correlates of finger-tapping task variations: an ALE meta-analysis NeuroImage. 2008;42:343–356.

[6] Turesky Ted K., Olulade Olumide A., Luetje Megan M., Eden Guinevere F. An fMRI study of finger tapping in children and adults Human Brain Mapping. 2018;39:3203–3215.

[7] Wüthrich Florian, Lefebvre Stephanie, Nadesalingam Niluja, et al. Test–retest reliability of a finger-tapping fMRI task in a healthy population European Journal of Neuroscience. 2023;57:78–90.

[8] Friston K. J., Holmes A. P., Worsley K. J., Poline J.-P., Frith C. D., Frackowiak R. S. J. Statistical parametric maps in functional imaging: A general linear approach Human Brain Mapping. 1994;2:189–210.

[9] Poline Jean-Baptiste, Brett Matthew. The general linear model and fMRI: Does love last forever? NeuroImage. 2012;62:871–880.

[10] Leibovici Didier G, Smith Stephen. Comparing groups of subjects in fMRI studies: a review of the GLM approach. FMRIB Technical Report. 2001.

[11] Monti Martin M. Statistical Analysis of fMRI Time-Series: A Critical Review of the GLM Approach Frontiers in Human Neuroscience. 2011;5.

[12] Zang Yufeng, Jiang Tianzi, Lu Yingli, He Yong, Tian Lixia. Regional homogeneity approach to fMRI data analysis NeuroImage. 2004;22:394–400.

[13] Lv Yating, Margulies Daniel S., Villringer Arno, Zang Yu-Feng. Effects of Finger Tapping Frequency on Regional Homogeneity of Sensorimotor Cortex PLOS ONE. 2013;8:e64115. Publisher: Public Library of Science.

[14] Farrugia Christine, Galdi Paola, Irazu Irati Arenzana, Scerri Kenneth, Bajada Claude J. Local gradient analysis of human brain function using the Vogt-Bailey Index Brain Structure and Function. 2024;229:497–512.

[15] Galea Kristian. Investigation of the effect of different functional MRI sequences on local homogeneity measures: implications for neuroimaging studies Master’s thesisUniversity of Malta 2024.

[16] Triantafyllou C., Hoge R. D., Krueger G., et al. Comparison of physiological noise at 1.5 T, 3 T and 7 T and optimization of fMRI acquisition parameters NeuroImage. 2005;26:243–250.

[17] Peirce Jonathan, Gray Jeremy R., Simpson Sol, et al. PsychoPy2: Experiments in behavior made easy Behavior Research Methods. 2019;51:195–203.

[18] Gorgolewski Krzysztof J., Auer Tibor, Calhoun Vince D., et al. The brain imaging data structure, a format for organizing and describing outputs of neuroimaging experiments Scientific Data. 2016;3:160044. Publisher: Nature Publishing Group.

[19] Esteban Oscar, Markiewicz Christopher J., Blair Ross W., et al. fMRIPrep: a robust preprocessing pipeline for functional MRI Nature Methods. 2019;16:111–116.

[20] Esteban Oscar, Ciric Rastko, Finc Karolina, et al. Analysis of task-based functional MRI data preprocessed with fMRIPrep Nature protocols. 2020;15:2186–2202.

[21] Woolrich Mark W., Jbabdi Saad, Patenaude Brian, et al. Bayesian analysis of neuroimaging data in FSL NeuroImage. 2009;45:S173–S186.

[22] Smith Stephen M., Jenkinson Mark, Woolrich Mark W., et al. Advances in functional and structural MR image analysis and implementation as FSL NeuroImage. 2004;23 Suppl 1:S208–219.

[23] Jenkinson Mark, Beckmann Christian F., Behrens Timothy E. J., Woolrich Mark W., Smith Stephen M. FSL NeuroImage. 2012;62:782–790.

[24] Tustison Nicholas J., Avants Brian B., Cook Philip A., et al. N4ITK: improved N3 bias correction IEEE transactions on medical imaging. 2010;29:1310–1320.

[25] Avants Brian B., Tustison Nicholas J., Song Gang, Cook Philip A., Klein Arno, Gee James C. A reproducible evaluation of ANTs similarity metric performance in brain image registration NeuroImage. 2011;54:2033–2044.

[26] Fischl Bruce. FreeSurfer NeuroImage. 2012;62:774–781.

[27] Klein Arno, Ghosh Satrajit S., Bao Forrest S., et al. Mindboggling morphometry of human brains PLOS Computational Biology. 2017;13:e1005350. Publisher: Public Library of Science.

[28] Tustison Nicholas J., Cook Philip A., Holbrook Andrew J., et al. The ANTsX ecosystem for quantitative biological and medical imaging Scientific Reports. 2021;11:9068. Publisher: Nature Publishing Group.

[29] Ciric Rastko, Thompson William H., Lorenz Romy, et al. TemplateFlow: FAIR-sharing of multi-scale, multi-species brain models Nature Methods. 2022;19:1568–1571.

[30] Fonov VS, Evans AC, McKinstry RC, Almli CR, Collins DL. Unbiased nonlinear average age-appropriate brain templates from birth to adulthood NeuroImage. 2009;47:S102.

[31] Fonov Vladimir, Evans Alan C., Botteron Kelly, Almli C. Robert, McKinstry Robert C., Collins D. Louis. Unbiased average age-appropriate atlases for pediatric studies NeuroImage. 2011;54:313–327.

[32] Jenkinson Mark, Bannister Peter, Brady Michael, Smith Stephen. Improved Optimization for the Robust and Accurate Linear Registration and Motion Correction of Brain Images NeuroImage. 2002;17:825–841.

[33] Greve Douglas N., Fischl Bruce. Accurate and robust brain image alignment using boundary-based registration NeuroImage. 2009;48:63–72.

[34] Power Jonathan D., Barnes Kelly A., Snyder Abraham Z., Schlaggar Bradley L., Petersen Steven E. Spurious but systematic correlations in functional connectivity MRI networks arise from subject motion NeuroImage. 2012;59:2142–2154.

[35] Behzadi Yashar, Restom Khaled, Liau Joy, Liu Thomas T. A component based noise correction method (CompCor) for BOLD and perfusion based fMRI NeuroImage. 2007;37:90–101.

[36] Patriat Rémi, Reynolds Richard C., Birn Rasmus M. An improved model of motion-related signal changes in fMRI NeuroImage. 2017;144:74–82.

[37] Alahmadi Adnan A. S. Effects of different smoothing on global and regional resting functional connectivity Neuroradiology. 2021;63:99–109.

[38] Weibull A., Gustavsson H., Mattsson S., Svensson J. Investigation of spatial resolution, partial volume effects and smoothing in functional MRI using artificial 3D time series NeuroImage. 2008;41:346–353.

[39] Glover Gary H. Deconvolution of Impulse Response in Event-Related BOLD fMRI1 NeuroImage. 1999;9:416–429.

[40] Parkes Linden, Fulcher Ben, Yücel Murat, Fornito Alex. An evaluation of the efficacy, reliability, and sensitivity of motion correction strategies for resting-state functional MRI NeuroImage. 2018;171:415–436.

[41] Satterthwaite Theodore D., Elliott Mark A., Gerraty Raphael T., et al. An improved framework for confound regression and filtering for control of motion artifact in the preprocessing of resting-state functional connectivity data NeuroImage. 2013;64:240–256.

[42] Winkler Anderson M, Ridgway Gerard R, Webster Matthew A, Smith Stephen M, Nichols Thomas E. Permutation inference for the general linear model Neuroimage. 2014;92:381–397.

[43] Kliemann Dorit, Adolphs Ralph, Armstrong Tim, et al. Caltech Conte Center, a multimodal data resource for exploring social cognition and decision-making Scientific Data. 2022;9:138.

[44] Bach Patrick, Hill Holger, Reinhard Iris, Gädeke Theresa, Kiefer Falk, Leménager Tagrid. Reliability of the fMRI-based assessment of self-evaluation in individuals with internet gaming disorder European Archives of Psychiatry and Clinical Neuroscience. 2022;272:1119–1134.

[45] Alarcon Nefi. Similarity in graphs: Jaccard versus the Overlap Coefficient 2019.

[46] Wilson Stephen M., Bautista Alexa, Yen Melodie, Lauderdale Stefanie, Eriksson Dana K. Validity and reliability of four language mapping paradigms NeuroImage: Clinical. 2017;16:399–408.

[47] Pimentel Marco A. F., Vilela Pedro, Sousa Inês, Figueiredo Patrícia. Localization of the hand motor area by arterial spin labeling and blood oxygen level-dependent functional magnetic resonance imaging Human Brain Mapping. 2011;34:96–108.

[48] Schotten Michel, Urbanski Marika, Valabregue Romain, Bayle Dimitri J., Volle Emmanuelle. Subdivision of the occipital lobes: An anatomical and functional MRI connectivity study Cortex. 2014;56:121–137.

