## Supplementary figures and images for "Testing the Vogt-Bailey Index using task-based fMRI across pulse sequence protocols"

### Supplementary_material

**VB Method 1**
**Sequence protocol: $2^3$ mm$^3$**

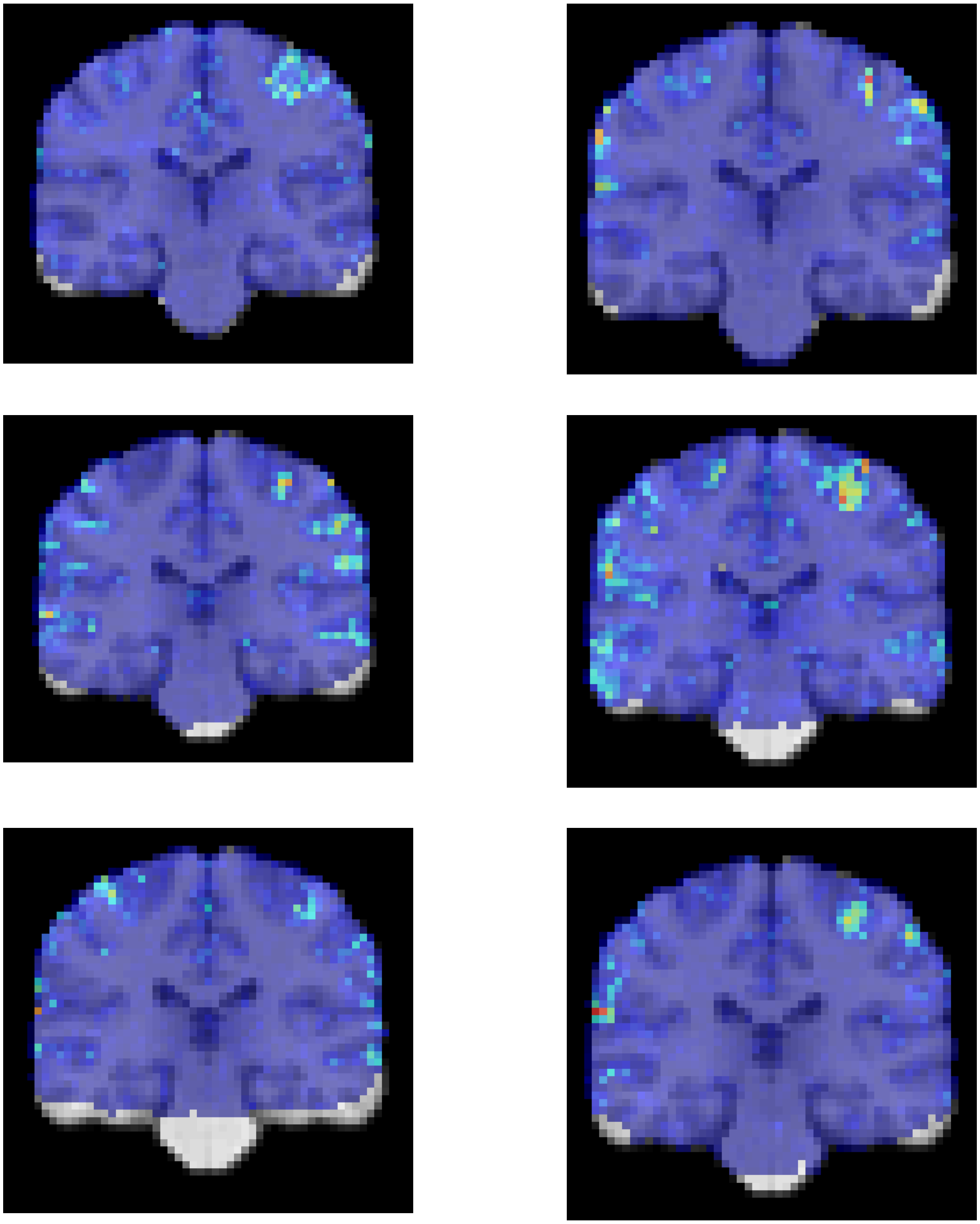

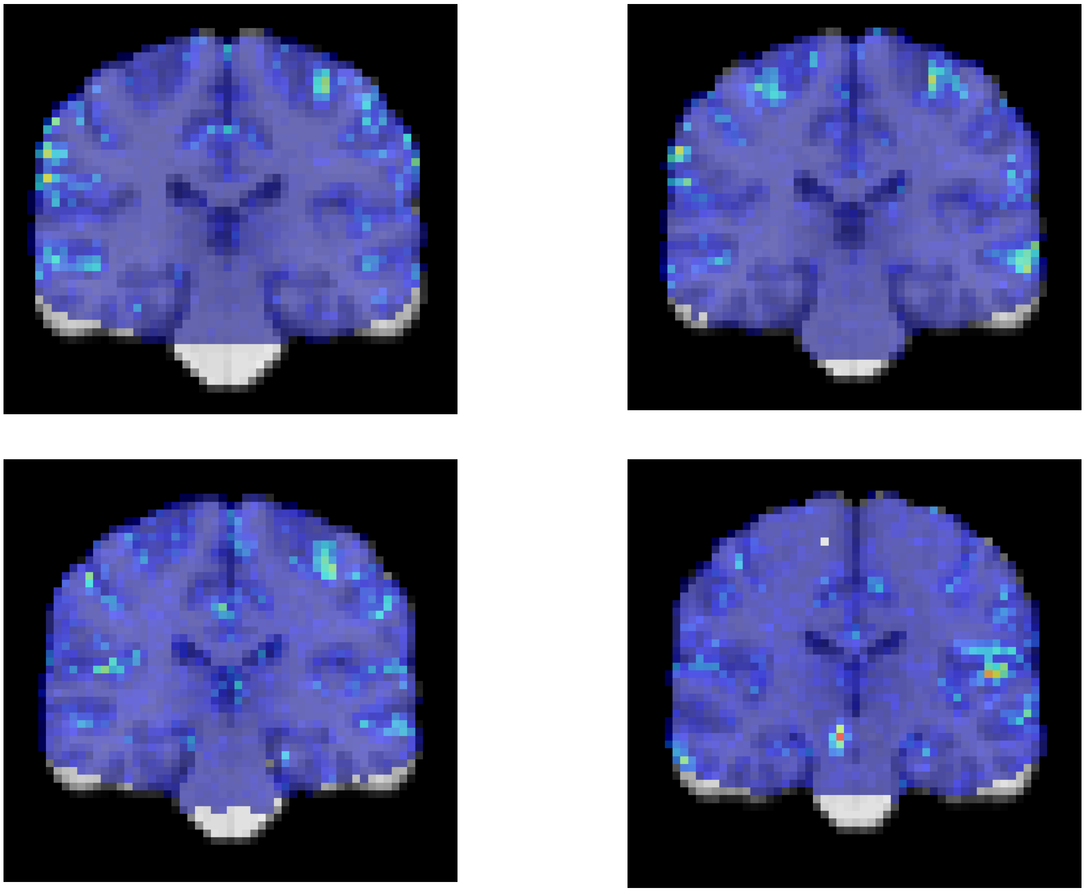

**VB Method 2**
**Sequence protocol: $2^3$ mm$^3$**

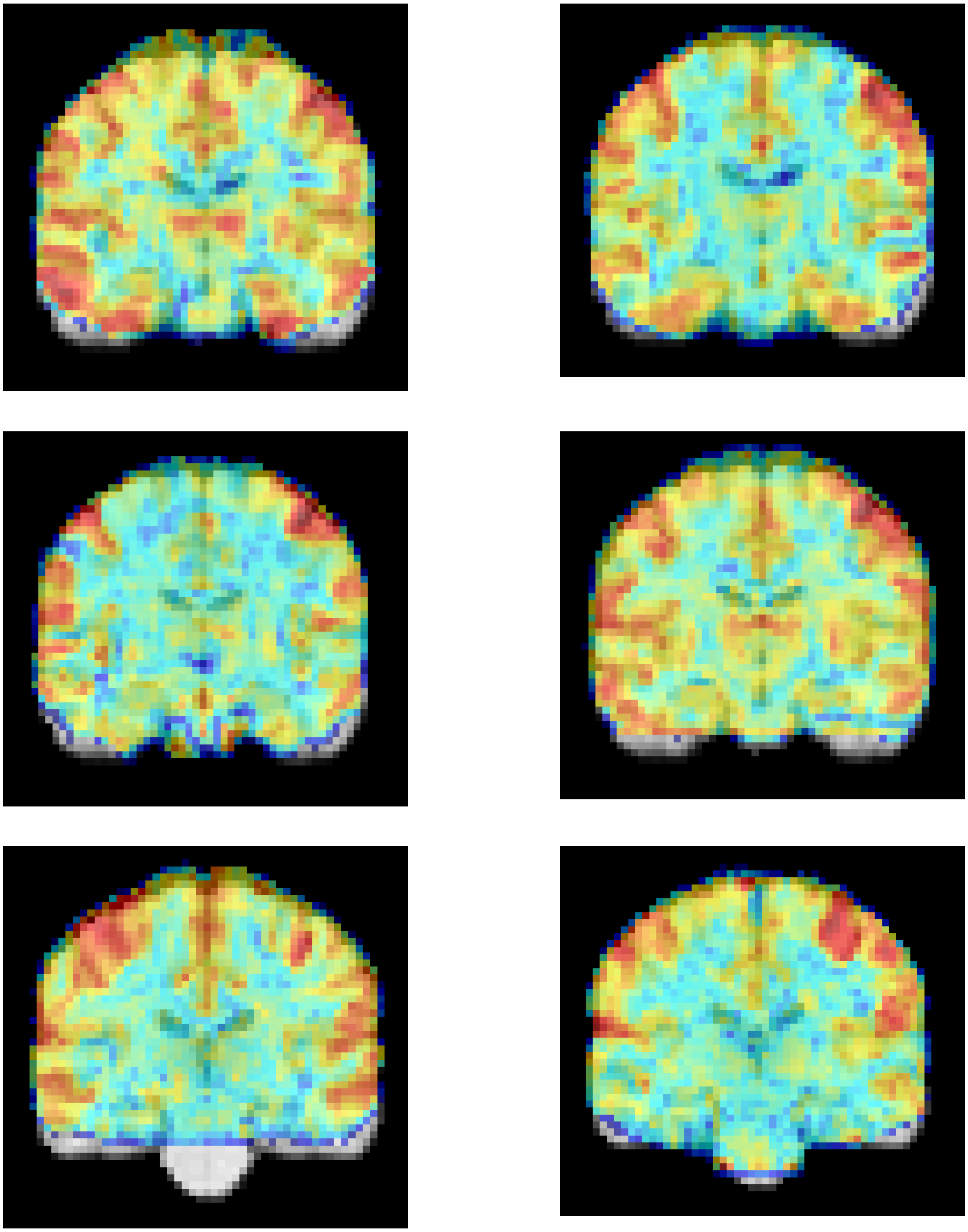

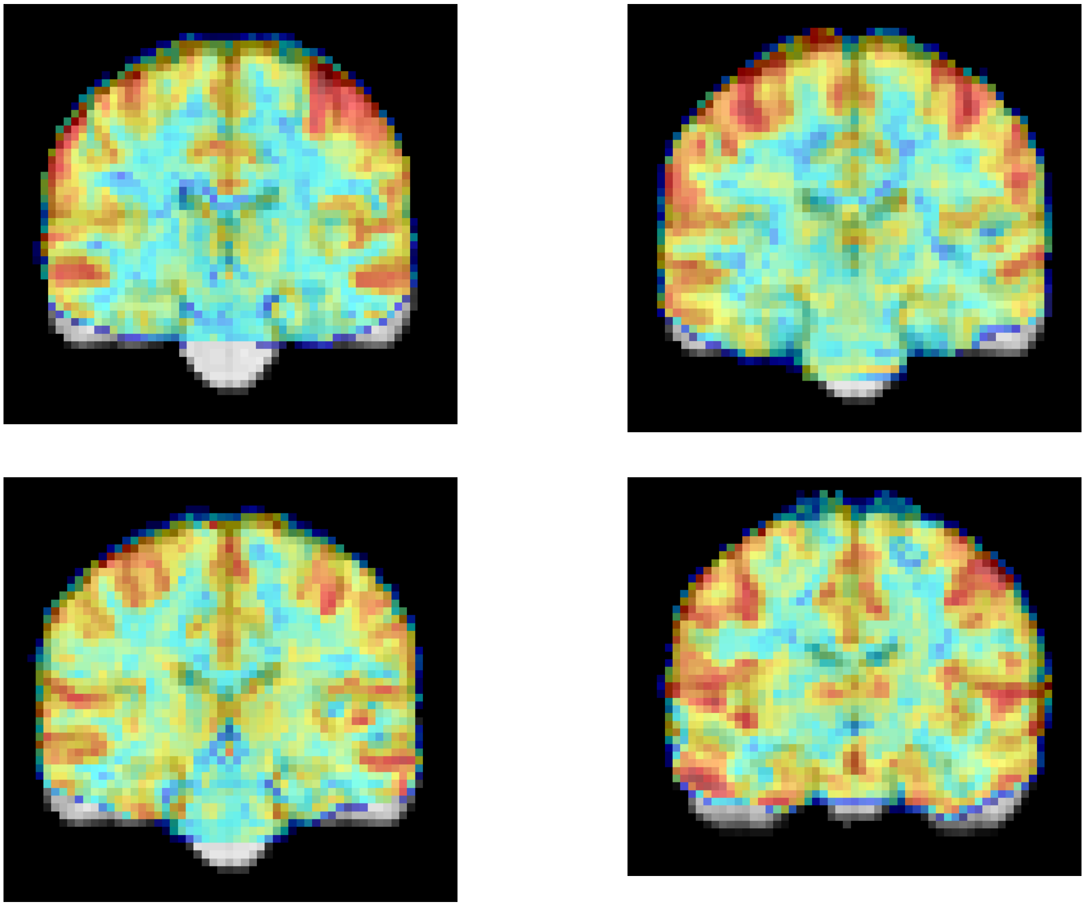
